# Universal toolset for mass spectrometric analysis of intracellular peptidome and small protein fraction

**DOI:** 10.1101/2024.12.06.627199

**Authors:** Sachin Kote, Artur Piróg, Jakub Faktor, Alicja Dziadosz, Natalia Marek-Trzonkowska

## Abstract

The analysis of native intracellular peptidome has gained significant attention in recent years. However, there is still a need for more knowledge regarding various sample preparation methods that facilitate efficient and reproducible recovery of peptides, which can then be analyzed using quantitative liquid chromatography-mass spectrometry. A similar situation exists in small proteome research, typically defined as polypeptides with masses of less than 100 amino acids, often too long for easy identification without enzymatic digestion. In this context, we describe a set of methods that involve simple denaturation and solid-phase extraction of polypeptides, applicable for isolating short intracellular polypeptides within the desired length range. Our work demonstrates the efficiency and reproducibility of these methods for quantitative analysis of the peptidome in mammalian cells. Additionally, we investigated the flexibility of adjusting the mass range through ultrafiltration. We have shown that these methods can be adapted for highly efficient enrichment and fractionation of small proteins, resulting in polypeptide isolates suitable for tryptic digestion and intact protein analysis. Moreover, we describe the use of freely available computational tools that can effectively manage the analysis of the resulting data. The research presented here will benefit the global scientific community in both fundamental (protein turnover, proteolytic processing, non-canonical open reading frames, *etc*.) and applied sciences (bioactive/neuro peptide discovery, precision medicine, vaccines, *etc*.), and other areas that could benefit from selective analysis of short native polypeptides.

## Introduction

The analysis of peptidomes has been constantly developing in recent years, expanding its applications to various biological sources. Most notably, peptidomics-based methods concentrate on analyzing biomarkers in body fluids, signaling peptides^1,2^, major histocompatibility complex (MHC)^3^ peptide ligands, or identification of proteins derived from short open reading frames (sORFs)^4,5^. We describe a novel framework for qualitative and quantitative analysis of native intracellular peptides and short proteins using mass spectrometry. The development of mass spectrometry instrumentation has substantially improved the identification of long and unspecific peptides, the availability of large sequence databases, and *de novo* peptide sequencing from mass spectra. Nevertheless, one of the most critical parts of any analysis is a reliable and reproducible sample preparation method, preferably enabling quantitative analysis.

Numerous intracellular peptides have been characterized in terms of their potential activity^6^. Nevertheless, research explicitly concentrating on the short peptide content of a cell’s interior is small. An insight into published reports suggests that current peptide extraction methods are often inefficient and require high input material. As shown in Table 1, identification depth varies significantly, from less than 1000 to more than 10000 peptide species, even from similar sample types. Input sample amounts differ by at least two orders of magnitude. Of note, the peptidomic studies listed in Table 1 are primarily semiquantitative, as the reported significance relies on uncorrected p-values. However, several truly quantitative experiments were also reported (e.g. Morgan et al. 2024, brain peptidome study^7^). Most methods rely on techniques selectively precipitating large proteins, combined with further clean-up steps, such as ultrafiltration and reverse-phase solid-phase extraction. For analysis of small proteome fractions, a wide range of methods, usually described as small protein enrichment is available^8,9^. They usually rely on size-based fractionation by ultrafiltration, SDS-PAGE, or selective precipitation. Table 1 suggests that most studies rely on multiple approach strategies to enhance the identification depth.

**Table 1.**
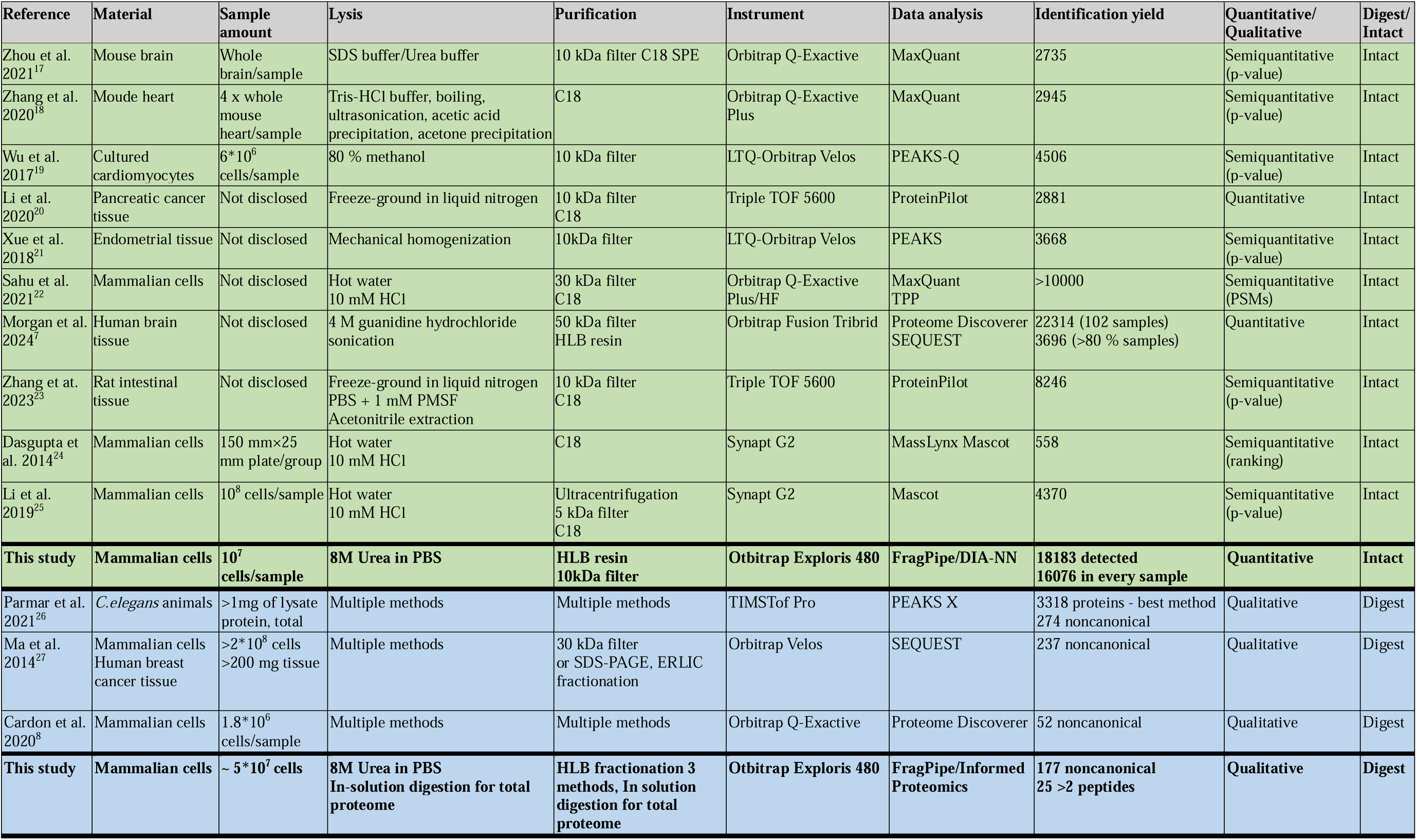
Comparison of methods and results of selected studies describing either peptidomics analyses performed on intact, short peptides (green) or small protein fraction studies performed via enzymatic digestion (blue).

**Table 2.** Sample lysis and peptide purification. PA method = pipet vigorously, 30 min vortex, 30 min sonification/RT, 20 min 17 kG centrifugation, add to supernatant 6 ml of 0.2% (FA). HLBF = HLB cartridge polypeptide eluate fractionation, HLBP = HLB cartridge polypeptide eluate purification for non-fractionated peptide eluates DIG = Digestion of fractions after initial analysis of intact peptides. A detailed description of the methods is provided in the supplementary material.

| Experiment | Cell line | Cell count | Cell lysis Buffer | Cell lysis Procedure | Fractionation | Peptide purification Procedure | Digestion |
| --- | --- | --- | --- | --- | --- | --- | --- |
| Cell lysis buffer comparison | A549 | 10<sup>7</sup> | 1 ml 8 M urea/PBS | PA | NO | HLBP 10 kDa cut-off filter | NO |
| Cell lysis buffer comparison | A549 | 10<sup>7</sup> | 200 µl 50% ACN, 0.1% FA | PA | NO | HLBP 10 kDa cut-off filter | NO |
| Cell lysis buffer comparison | A549 | 10<sup>7</sup> | 50 µl of 100% FA and 200 µl of H<sub>2</sub>O | PA | NO | HLBP 10 kDa cut-off filter | NO |
| Mass cut-off filter comparison | MG-132 treated A549 cells | 8x10<sup>6</sup> | 1 ml 8 M urea/PBS, 1 x Roche Protease Inhibitors | PA | NO | HLBP 3,10 and 30 kDa cut-off filter | NO |
| Polypeptide fractionation experiment | MG-132 treated A549 cells | 8x10<sup>6</sup> | 1 ml 8 M urea/PBS, 1mM PMSF and 1mM EDTA (omits peptide interference from PI) | PA | HLBF | HLBF itself | YES, after analysis of intact peptides (DIG) |
| Melanoma quantitative cellular peptidomic experiment | A375 | 10<sup>7</sup> | 1 ml 8 M urea/PBS, 1 x Roche Protease Inhibitors | PA | NO | HLBP 10 kDa cut-off filter | NO |
| Analysis of the effect of proteasome inhibition | SUP-T1 | 2x10<sup>7</sup> | 1 ml 8 M urea/PBS, 1mM PMSF and 1mM EDTA | PA | NO | HLBP 10 kDa cut-off filter | NO |

**Table 3.** Summary of liquid chromatography methods used in described experiments.

| LC method | Gradient length (min)/type | Initial/Final %B | Sample Loading length (min) | Column temperature (°C) | Trap column ID/L/particle size | Trap column packing | Analytical column ID/L/particle size (µm/mm/µm) | Analytical column packing |
| --- | --- | --- | --- | --- | --- | --- | --- | --- |
| 1 (Lysis buffer comparison) | 90/Linear | 2.5/40 | 10 | 35 | 0.3/5/5 | C18 PepMap 100 Å | 75/250/2 | C18 PepMap 100 Å |
| 2 (10kDa comparison) | 90/Linear | 2.5/40 | 10 | 50 | 0.3/5/5 | C18 PepMap 100 Å | 075/250/2 | C18 PepMap 100 Å |
| 3 (30kDa comparison) | 90/Linear | 2.5/40 | 10 | 60 | 0.3/5/5 | C18 PepMap 100 Å | 75/250/2 | C18 PepMap 100 Å |
| 4 (3kDa comparison) | 90/Non-linear | 2.5/20 (40min)/60 (50min) | 10 | 35 | 0.3/5/5 | C18 PepMap 100 Å | 75/250/2 | C18 PepMap 100 Å |
| 5 (Intact short) | 40/Linear | 2.5/40 | 5 | 50 | 0.3/5/5 | C18 PepMap 300 Å | 75/150/1.7 | BEH C18 300 Å |
| 6 (Intact long) | 90/Linear | 2.5/40 | 10 | 50 | 0.3/5/5 | C18 PepMap 300 Å | 75/150/1.7 | BEH C18 300 Å |
| 7 (Bortezomib) | 90/Linear | 2.5/40 | 10 | 35 | 0.3/5/5 | C18 PepMap 300 Å | 75/150/1.7 | BEH C18 300 Å |

**Table 4.**
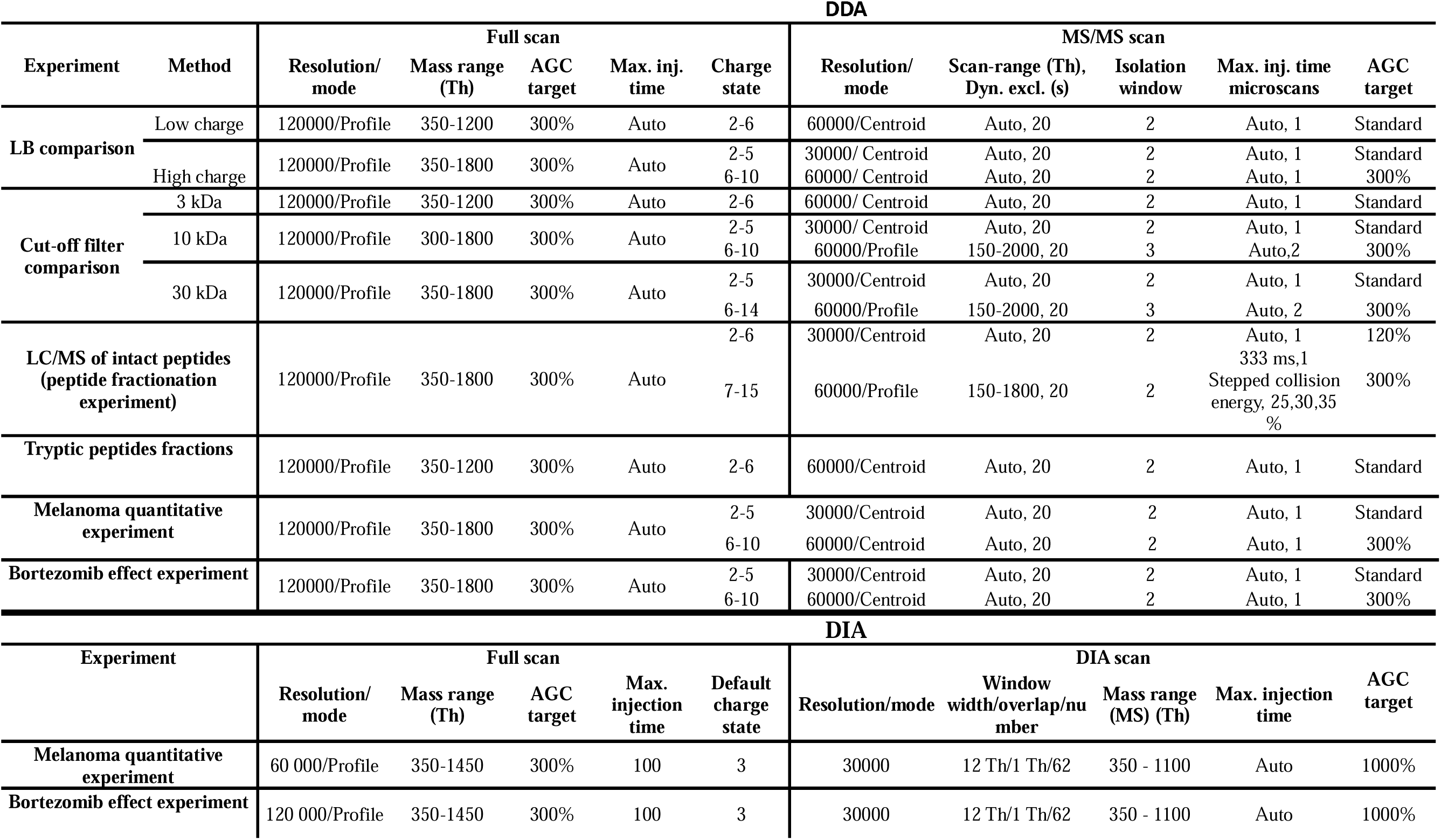
Summary of mass spectrometry methods used in described experiments.

Here, we have focused on developing a method to isolate and characterize the comprehensively intracellular peptidome and short proteome. Intracellular peptidome identification and quantification could be applied to unfold novel research directions (Figure 1). It may be used to discover bioactive peptides originating from protein degradation, non-canonical translation events, or translation of sORFs^10,11^. Alternatively, it can be applied to monitor cellular activities involving the production of peptides, such as protein degradation or pioneering translation events, which are further connected with other important processes such as nonsense-mediated decay (NMD) or antigenic peptide presentation in context of MHC molecules^12,13^. Intracellular peptides can be identified in cultured cells and tissue samples^14,15^. However, the isolation and analysis of the tissue intracellular peptides are challenging due to extensive contamination derived from the extracellular matrix.

**Figure 1.**
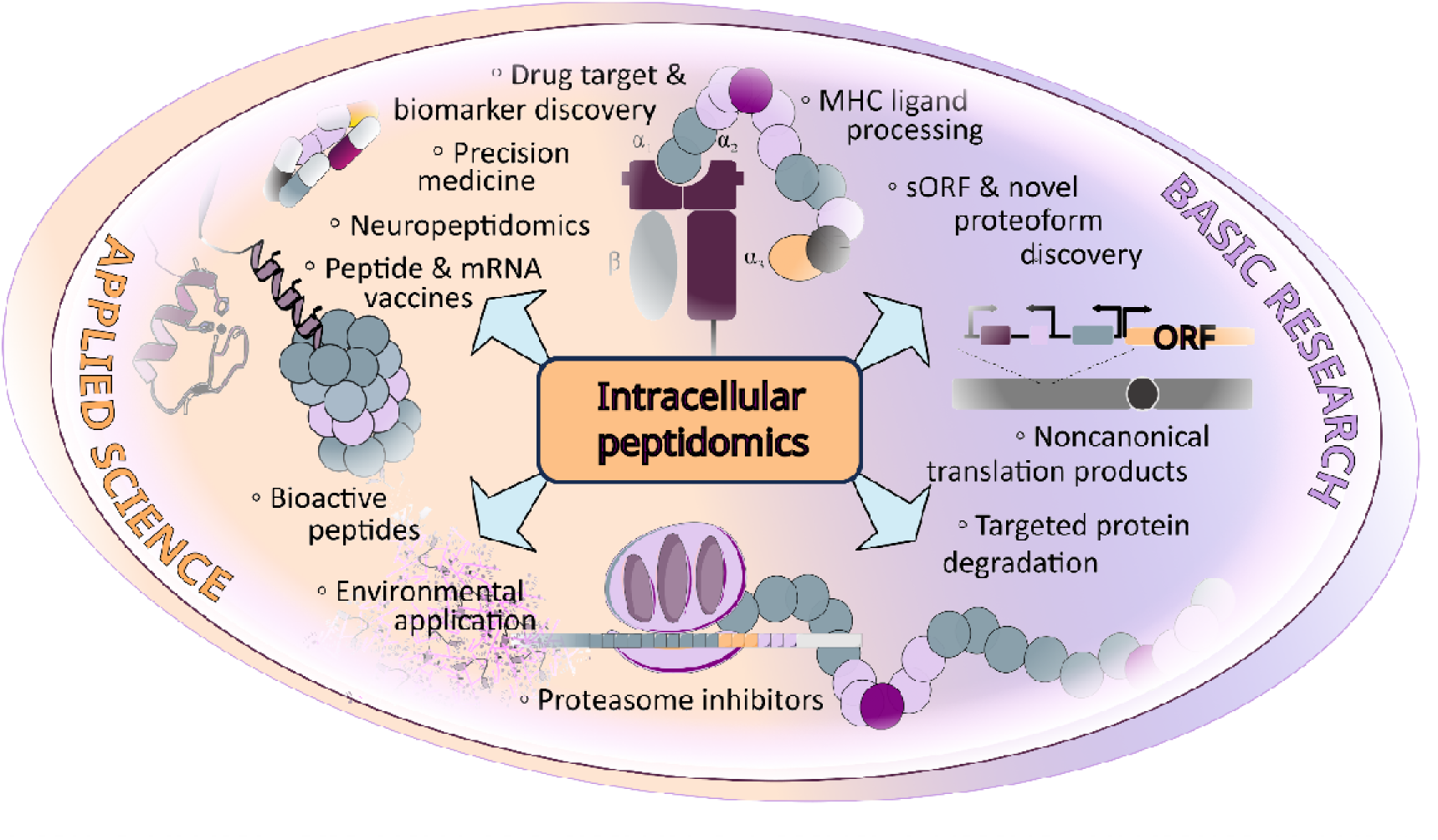
Potential applications of intracellular peptidome analysis.

**Figure 2.**
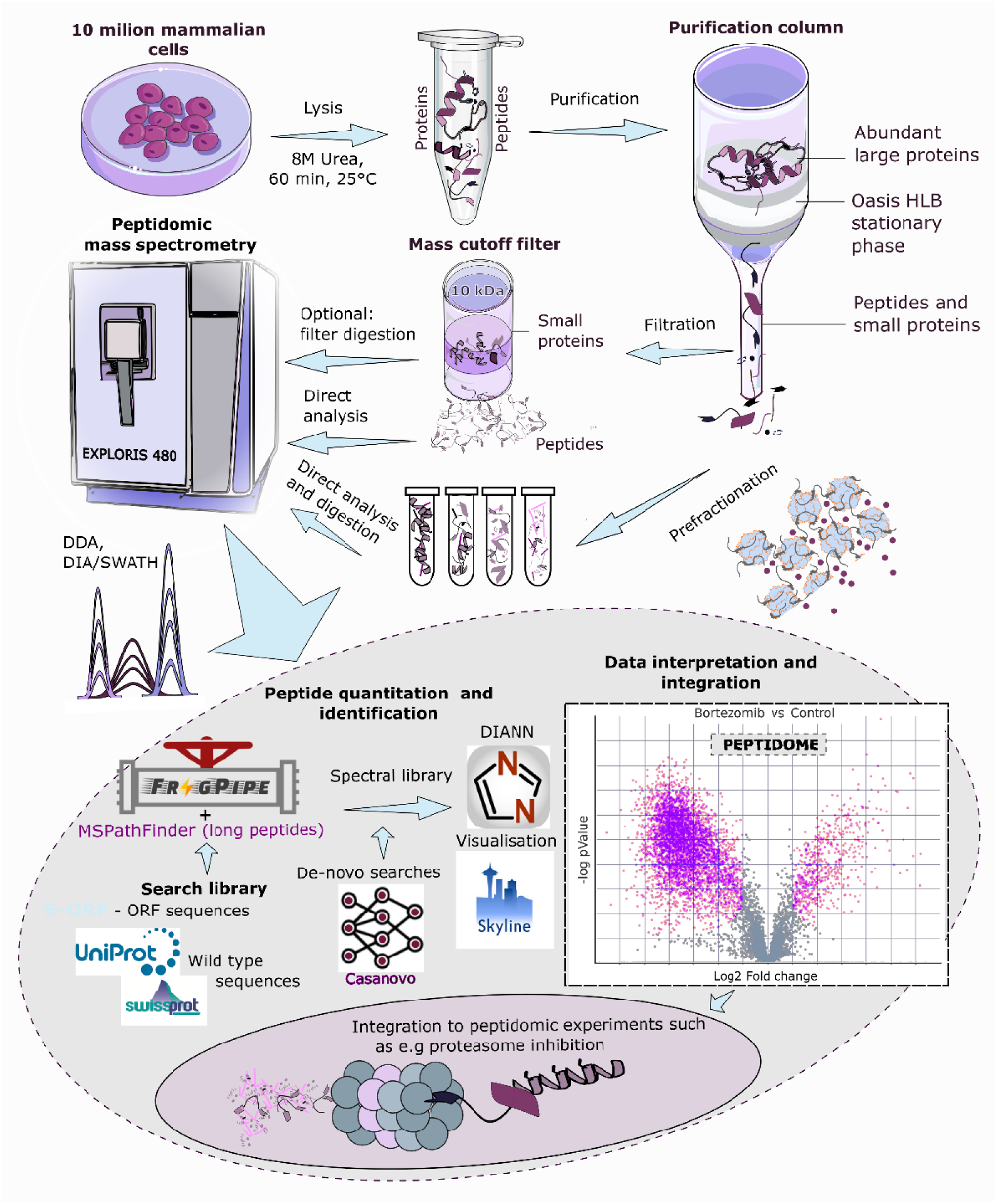
Overview of novel sample preparation methods, data acquisition, and data analysis for intracellular peptidome experiments.

The current study presents a novel, reproducible, and efficient method for extracting and quantitatively analyzing intracellular peptides. The approach is based on initial SPE sample cleaning by HLB resin^16^, accompanied with ultrafiltration to isolate peptides with a defined length range. The method variations described in the manuscript are designed to provide efficient analysis of intracellular peptides. Our method works with a limited sample amount ranging from one 8 cm plate with 10^7^ cells, providing sufficient mass spectrometry data for deep quantitative peptidome analysis. In addition, modern mass spectrometry instrumentation and advanced data analysis techniques allowed us to perform a successful in-depth identification and data-independent acquisition (DIA) quantification of 6000 to 16000 intracellular peptides. Moreover, this method was modified for small protein enrichment, providing excellent recovery and easy protein-level sample fractionation.

## Results

### Lysis buffer composition and filter size affect the physiochemical properties of isolated peptide population

Initially, inspired by existing literature, we screened several lysis buffers, including organic solvent-based (50% acetonitrile in 0.1% formic acid - ACN), acid-based (100 % / 20 % formic acid - FA), and chaotropic-based (8 M buffered urea - Urea) buffers to lyse the cells. Following, lysates were diluted 6 to 30 times before solid-phase extraction, leading to greater lysis methods’ flexibility. This approach reduced the concentration of compounds interfering with resin binding, such as urea or organic solvents, before sample loading to the OASIS column.

As shown in Figure 3, urea lysis outperforms acetonitrile and formic acid-based methods regarding the number of identified peptides. Nevertheless, many peptides remain unique to ACN and FA methods. To check whether this difference comes from distinct properties of peptides, we have compared their physiochemical properties. Acetonitrile and formic acid lysis solutions tend to isolate more hydrophilic and acidic peptides than urea. This observation should be considered when investigating a specified type of intracellular peptide. In addition, various lysis methods can be used in parallel to increase the number of identifications while aiming to isolate a broad range of intracellular peptides. This might be of great importance for screening/discovery studies.

**Figure 3.**
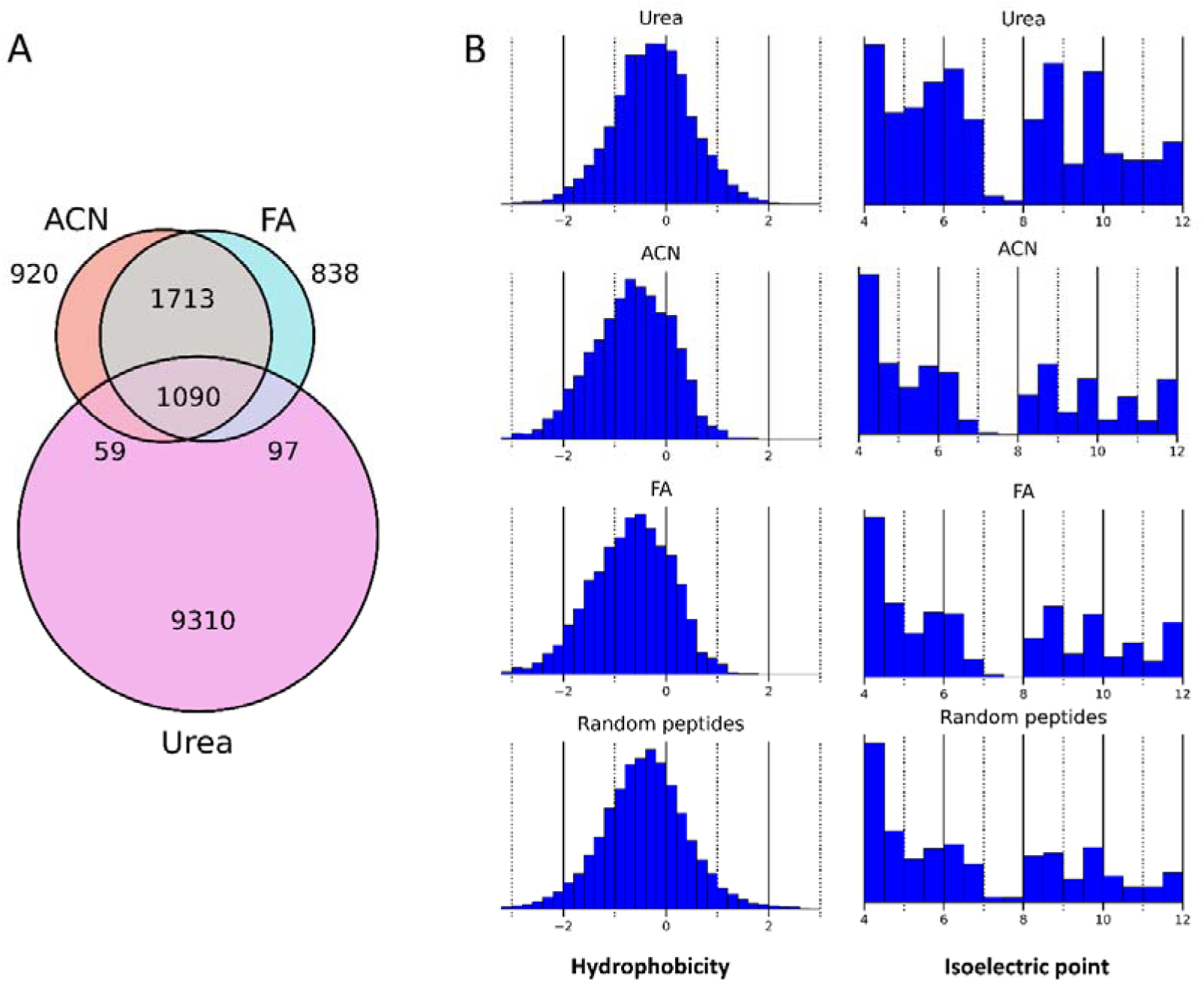
Comparison of different methods of cell lysis for intracellular peptidome isolation. A) Venn diagram showing overlap of peptide populations detected in samples obtained using the following lysis solutions: acetonitrile in formic acid (ACN), formic acid (FA) and buffered urea (Urea). B) Comparison of peptide hydrophobicity and isoelectric point distributions of peptides detected in samples obtained using different lysis methods (ACN, FA and Urea). The peptide reference was created as the population of peptides randomly selected from all human proteins, preserving the length distribution of peptides detected in all samples.

Next, we aimed to reduce the presence of very long peptides in isolates by optimizing the ultrafiltration step. We have compared the results of intracellular peptidome ultrafiltration after implementing filters with various mass cut-off values, particularly 3, 10 and 30 kDa. It is important to note that mass cut-off values are approximate, and these filters are rather designed to trap the concentrated sample on the filter top than provide a filtrate with a particular mass cut-off. Therefore, obtained results do not directly correspond to the filter mass cut-off value and may vary depending on the exact conditions of filtration. Figure 4A shows deconvoluted retention time/mass maps of representative samples prepared using 3, 10 and 30 kDa mass cut-off filters (non-deconvoluted counterparts are shown in Supplementary Figure 1). Figure 4B summarizes the total intensity of detected peptide features in these filtrates. As expected, the lowest peptide yield was obtained after using 3 kDa mass cut-off filter. However, the application of this filter is entirely justified when low molecular weight peptides are studied. In general, 10 kDa filter provides the best balance between recovery and reduction of high molecular weight peptides, which are less compatible with LC/MS analyses. We want to highlight that the actual peptide mass distribution might diverge from the distribution of detected peptide signals, as the efficacy of LC separation and ionization can significantly differ for short and long peptides. Nevertheless, the trend is expected to be similar, regardless of the influence of separation and ionization efficiency. This experiment offers insights into the types of peptide mass distributions that can be identified by varying the molecular weight cut-off.

**Figure 4.**
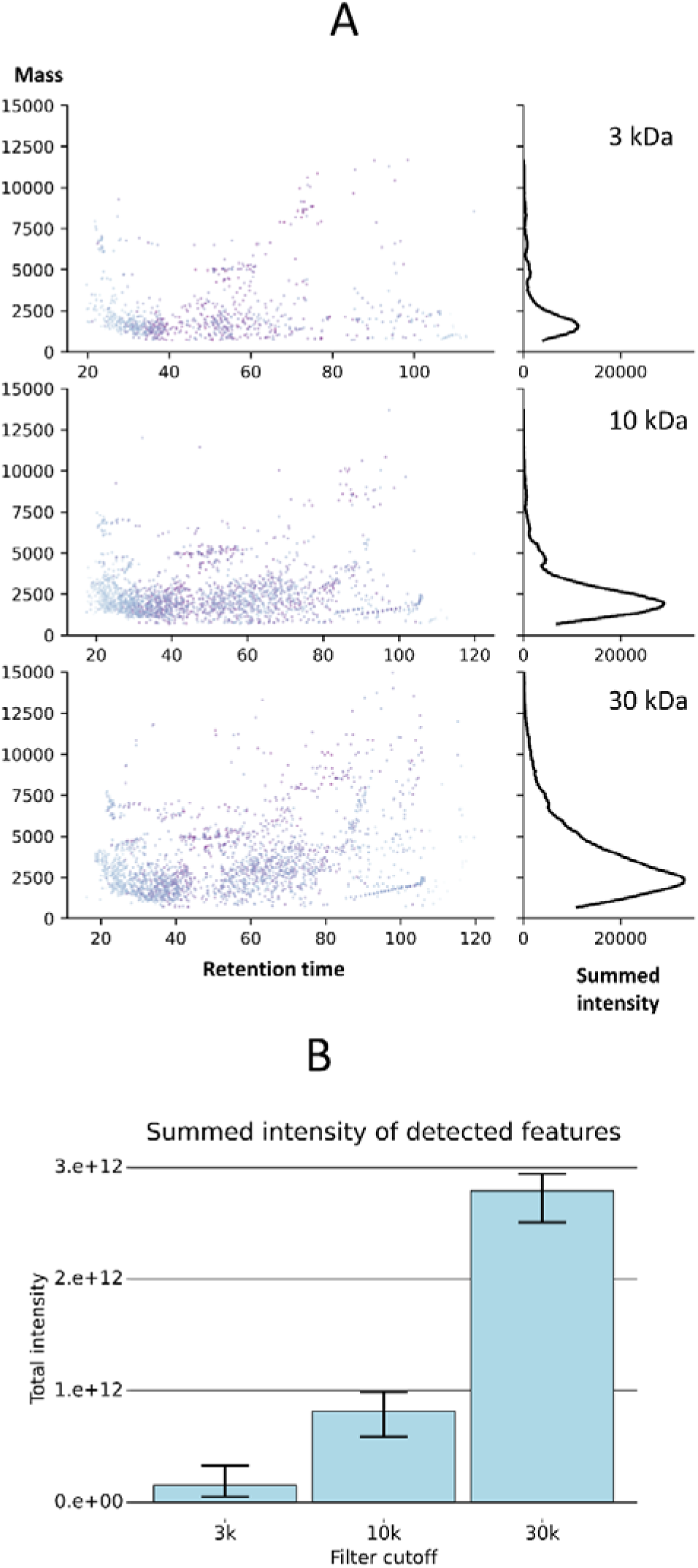
Comparison of results obtained with different mass cutoff filters. A) Detected deconvoluted peptide feature mass/retention time distributions of representative sample runs obtained using 3 kDa, 10 kDa and 30 kDa cut-off filters. The right panel histograms summarize overall intensity vs. mass distribution. B) Overall feature intensity sum for samples obtained using different filter sizes (n=3 per each filter type). Bar heights refer to mean; whiskers indicate minimum and maximum values.

### Optimized peptide extraction method enables precise quantitative studies of intracellular peptidome

Next, we checked if intracellular peptidomic filtrates prepared by the optimized method are compatible with the data-independent acquisition (DIA)-based peptide quantitation pipeline. Our objective was to ascertain if the optimal DIA method mass range would cover the broad mass and length range of peptide infiltrates. Hence, we directed our efforts towards determining whether there were any limitations on the maximum length of reproducibly quantifiable peptides. Fragmentation of wide mass windows results in complex fragmentation patterns and interferences between signals, a common limitation of DIA-based quantitation, even for short tryptic peptides.

We have tested our pipeline on intracellular peptidome filtrates prepared from wild-type and p53 knockout human melanoma cell lines (A375). We have chosen a 10 kDa filtrate for quantitative analysis, as the utilized filter mass cut-off omits longer peptide species while preserving a reasonable amount of material for analysis. We have used liquid chromatography (LC) separation conditions typical for peptide analysis (see details in the Methods section). Data-dependent acquisition (DDA) and DIA methods were slightly modified, utilizing higher resolution than typical proteomic experiments focused on tryptic peptide quantification^28^. Namely, DIA fragment spectra resolution was set to 30 000 and default charge state to 3+, enhancing more extended peptide characterization. The high-resolution acquisition, combined with a cycle time of ≈ 4.5 s on Orbitrap Exploris 480 (ThermoFisher), enabled sufficient points per peak (n∼10).

A wide range of non-specific intracellular peptide lengths was quantified successfully with the established protocols. Further, a comparison of A375 wild-type (WT) peptide detection efficiency and reproducibility in data-dependent acquisition (DDA) and data-independent acquisition (DIA) datasets are shown in Supplementary Figure 3.

Length distribution comparison of identified and quantified peptides without missing values suggests that longer peptide quantification is efficient. Our DIA method enabled the quantitation of 16076 intracellular peptides (Figure 5). We mapped the charge state of the peptides distribution of identified peptides from intracellular peptidome analysis, and a typical proteomic experiment was done on trypsin-digested samples, as shown in Supplementary Figure 4.

**Figure 5.**
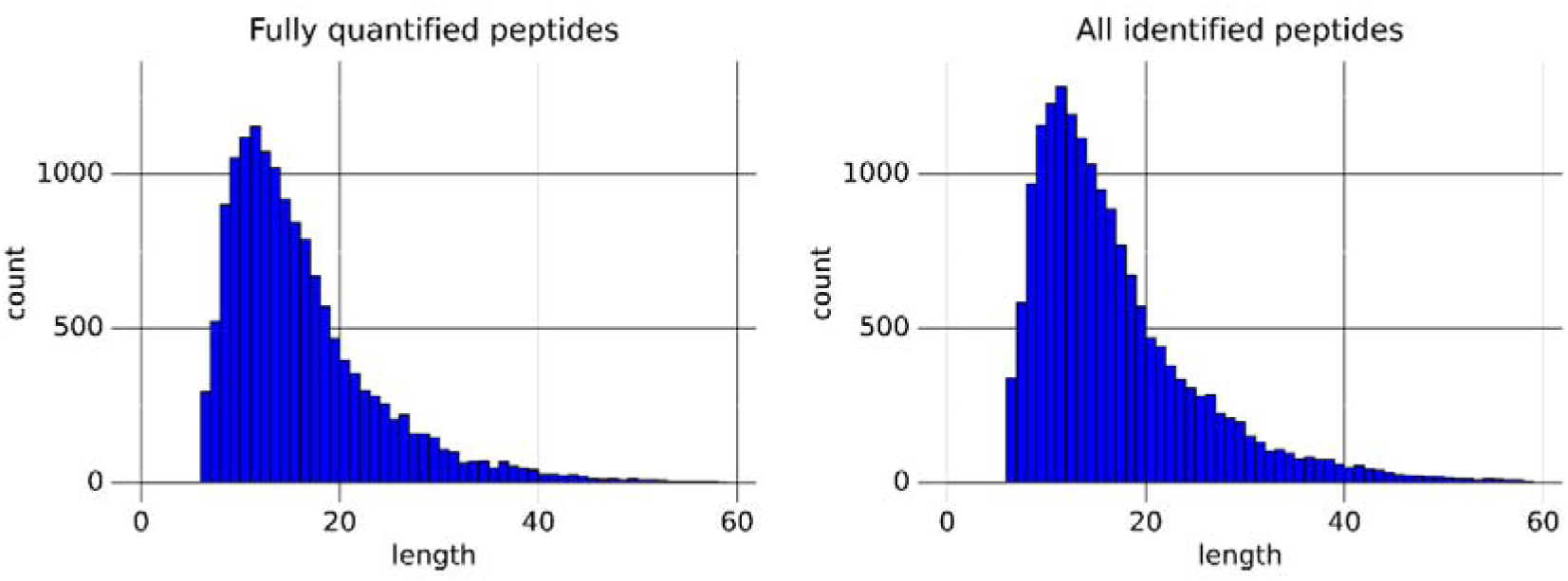
A length distribution histogram shows peptides quantified without any missing value (left) and all identified peptides (right) in a melanoma intracellular peptidomics experiment.

Next, we checked the reproducibility of quantitative analysis and its ability to detect significantly regulated intracellular peptides. Supplementary Figure 2 summarizes the quantification and detection reproducibility among three biological replicates of A375 wild-type and p53 knockout intracellular peptidome filtrates. We could maintain the intra-group correlation factor (Pearson R^2^) above 0.8. and to detect 74 regulated peptides (Figure 6) potentially originating from 37 proteins. A complete list of identified regulated peptides is listed in Supplementary Table 1.

**Figure 6.**
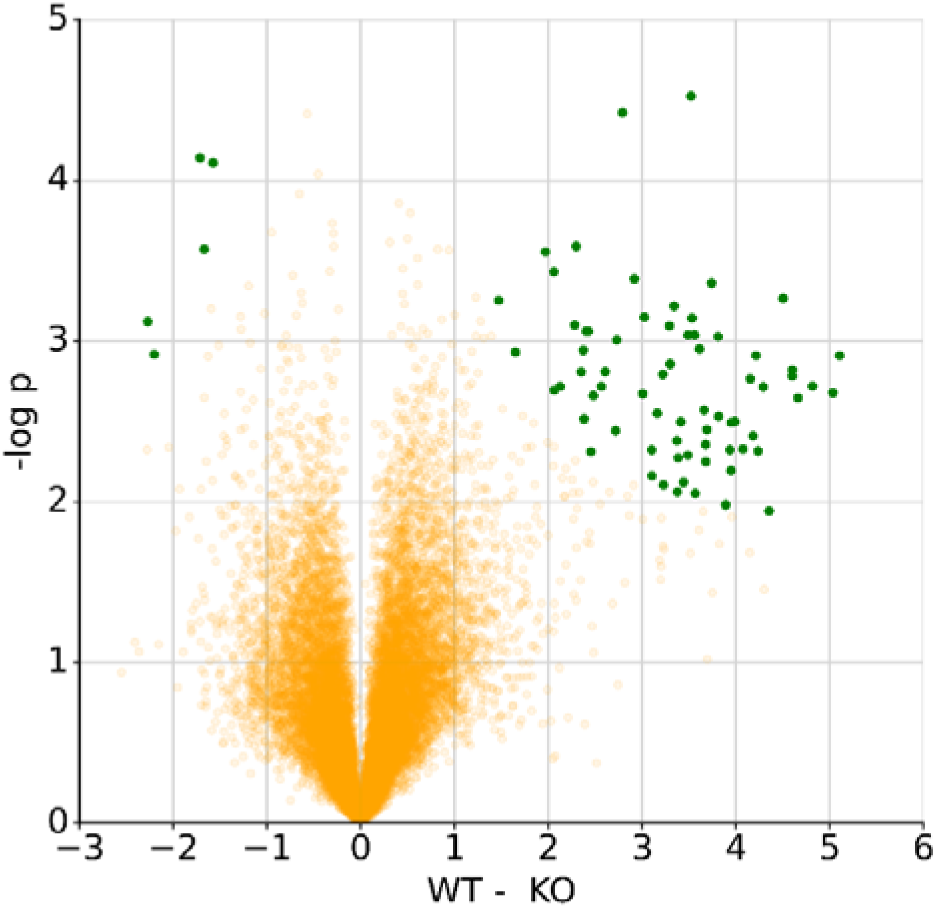
Overall distribution of peptidome differences between wild-type and p53 knockout A375 cell lines. The figure depicts a peptide-level volcano plot showing differences between peptidomes of wild-type and p53 knockout A375 cell lines. Green dots indicate peptides whose abundance differences are considered statistically significant.

The interpretation of biological meaning reflected in cellular peptidome changes remains challenging, as knowledge regarding its functional aspect is limited. Vast portion of intracellular peptides comes from full-length proteins. Therefore, many changes detected in intracellular peptidome reflect the extent of native protein proteolytic processing or changes in their expression. We performed a functional analysis demonstrating several examples of biologically meaningful results to prove the possible application of the newly developed quantitative intracellular peptidomics method. We have identified peptides originating from proteins related to the p53 pathway that were affected by P53 knockout (Figure 7). Sequestosome-1-derived peptides that show concerted downregulation (4-8 fold) in p53 KO cells represent the most notable changes. This protein functions as an adapter for autophagy, mainly of polyubiquitinated proteins^29^. In general, p53 is known to induce autophagy^30^, and it can be expected that protein amount and turnover reduction will result in a marked decrease in its probable degradation products. Another interesting observation is the presence of peptides derived from the C-terminal part of Y-box-binding protein 1 (YB-1). This protein is known to be proteolytically processed. Its C-terminal part is retained in cytoplasm, whereas the rest of the protein transfers to the nucleus. This process is at least partially stimulated by p53^31^. Variable changes in amounts of different peptides derived from YB-1 protein suggest modification of its proteolytic processing. Another known interactor of p53 is protein S100A4^32^. It plays an important, nevertheless not well-explained, role in the regulation of cell motility, survival, and metastasis^33,34^. Overall, the S100A4-derived peptide level is markedly reduced, and we have found that its C-terminal peptide was previously detected in another study^35^.

**Figure 7.**
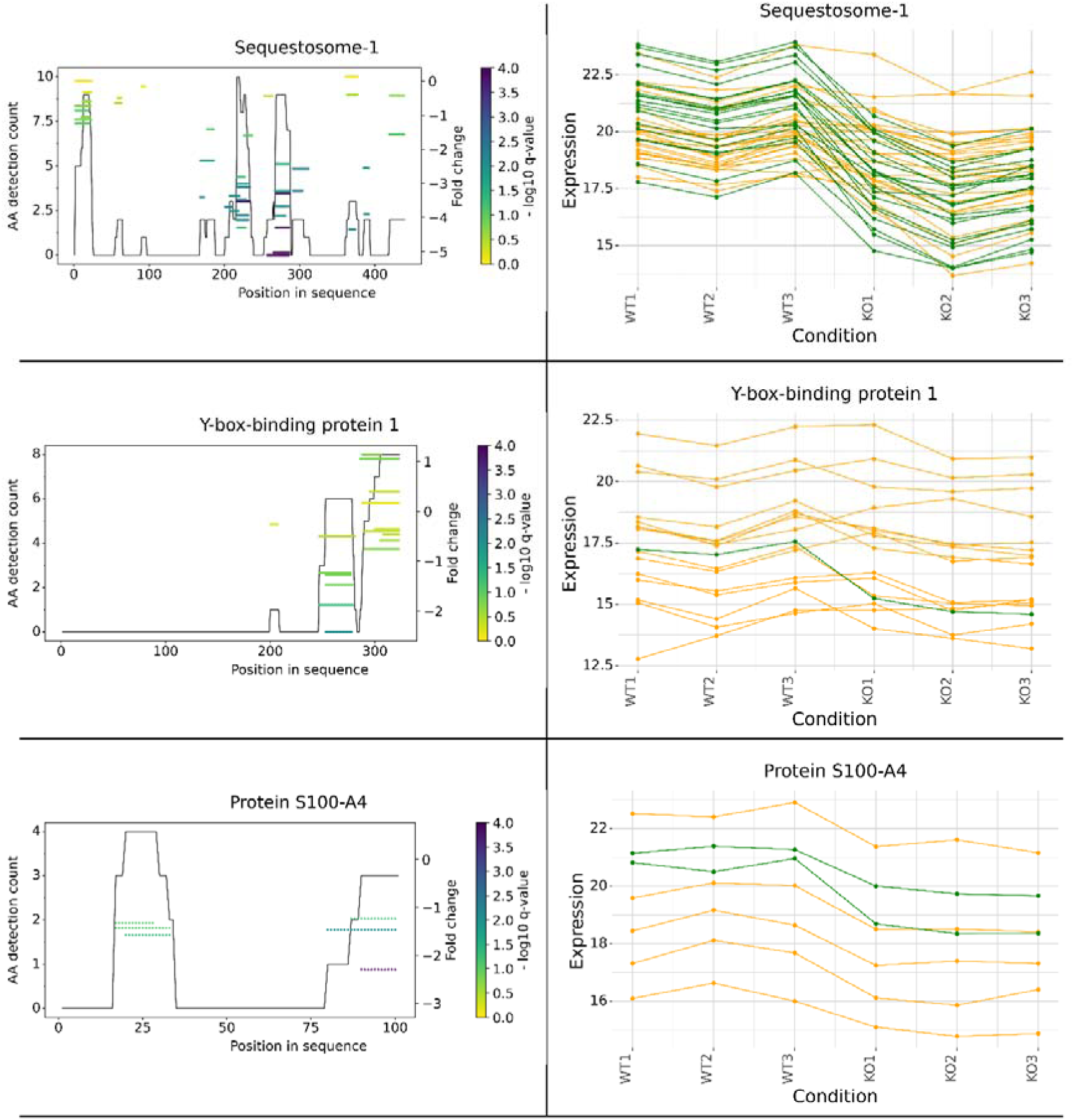
Peptide coverage and peptide fold change plots of selected proteins associated with p53 knockout. Left side graphs present detection coverage across the protein sequence as a black line. Horizontal lines symbolize individual detected and quantified peptides. The vertical position of the line indicates measured fold change, and color reflects the Welch test multiple test corrected p-value (q-value). The right-side graphs present peptide quantifications across the samples. The green color indicates that the peptide was detected as significant by the moderated Welch test performed in Perseus.

Furthermore, we attempted to characterize several aspects of obtained intracellular peptide population globally. As shown in Figure 8, peptide position analysis mapped into possible full-length proteins of origin exhibited a clear overrepresentation of N-term and C-terminal peptides. Similar observations have been published before; however, there is a high variability among the datasets, and the reason for the enrichment of C and N-term peptides in intracellular filtrates is not yet well described^36–38^.

**Figure 8.**
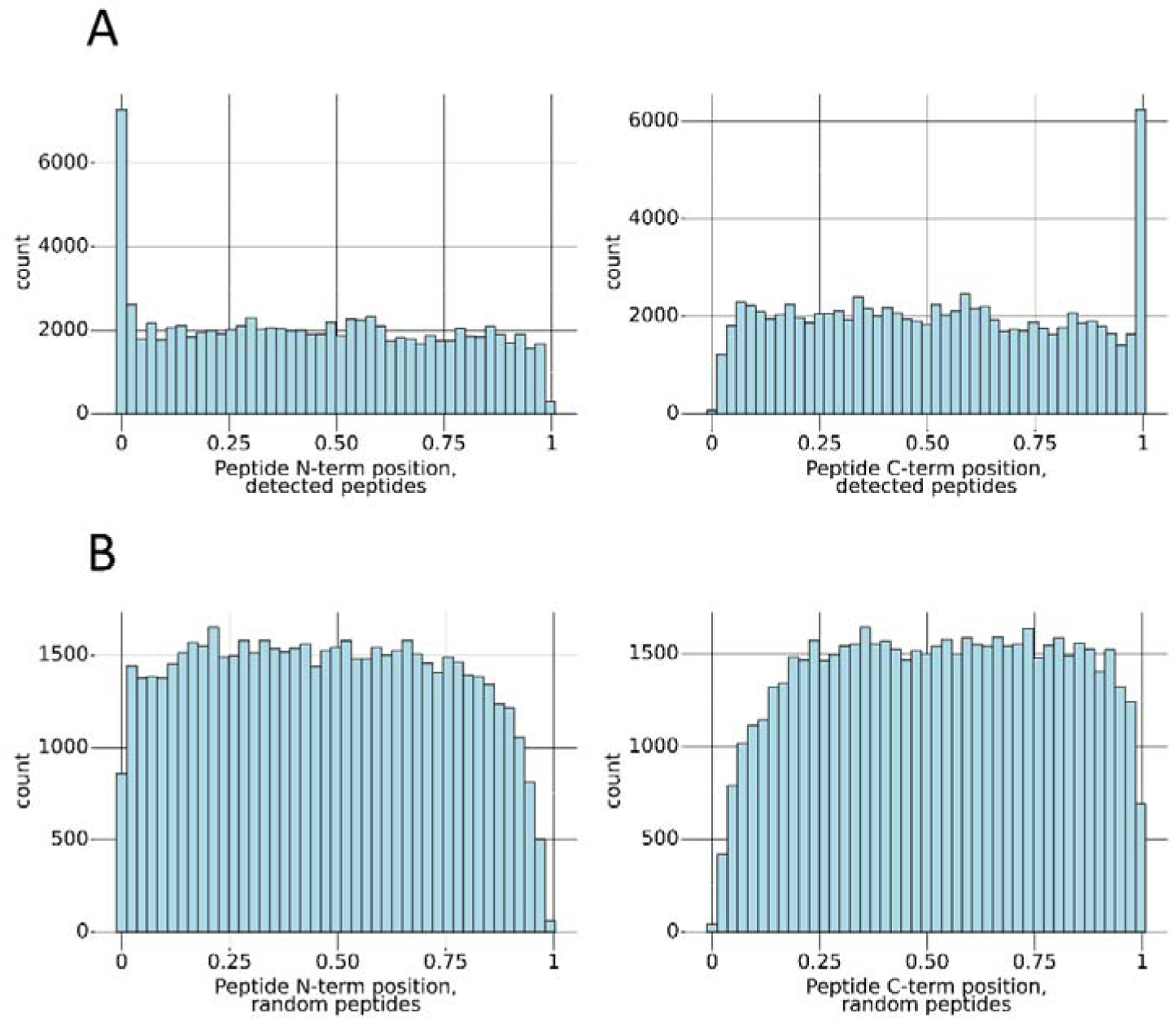
Intracellular peptides mapped into possible source protein sequences to detect peptide termini location in the A375 melanoma cell line. Distribution A shows the locations of termini of detected peptides. Distribution B shows the location of the termini in random peptides of the same length distribution.

The majority of intracellular peptides may come from the proteolysis of longer peptides. The lesser part is a result of the direct translation of short polypeptides. Therefore, amino acid composition analysis of peptide termini should reflect the diversity of protease specificities, mostly proteasomal subunits. We have compared the peptide flank region amino acid composition −3 and +3 amino acids downstream from N-term and upstream from C-term in identified peptides, respectively.

Figure 9 shows that cut sites at P1 position on both N and C terminus flanks are enriched in hydrophobic amino acids (L, F, M, Y, W), suggesting chymotryptic-like specificity of the protease involved in peptide processing. Strikingly, there is a strong enrichment for cysteine residue at position P-1 in the N-term flank, slight at C-term P1’ flank and a reduction of cysteine residues inside the peptide sequence. Calculating for whole length of the peptide, cysteines represent 1.8% of all amino acids in the peptide sequence of randomly generated peptides compared to 0.3% in peptides derived from the experimental dataset. It should be noted that the intracellular peptidomic dataset was generated without reduction and alkylation, which may explain the underrepresentation of cysteines in the identified peptides as they form disulfide bridges but does not explain overrepresentation at position P-1 on the N-term flank. A similar observation was previously reported in mouse brain peptidomic study^14^. Overall, terminus enrichment and flank amino acid composition suggest that a substantial part of these peptides do not come from direct proteasomal cleavage. Methods specifically targeting proteasomal peptides yielded a different distribution, better reflecting the proteasome subunit specificity and much lesser bias toward terminal peptides^39^. However, the authors also found surprisingly few peptides with tryptic-like termini. This effect may be cell-line specific, as in the experiment described below, which detected protease specificities that matched possible proteasome digestion patterns more closely.

**Figure 9.**
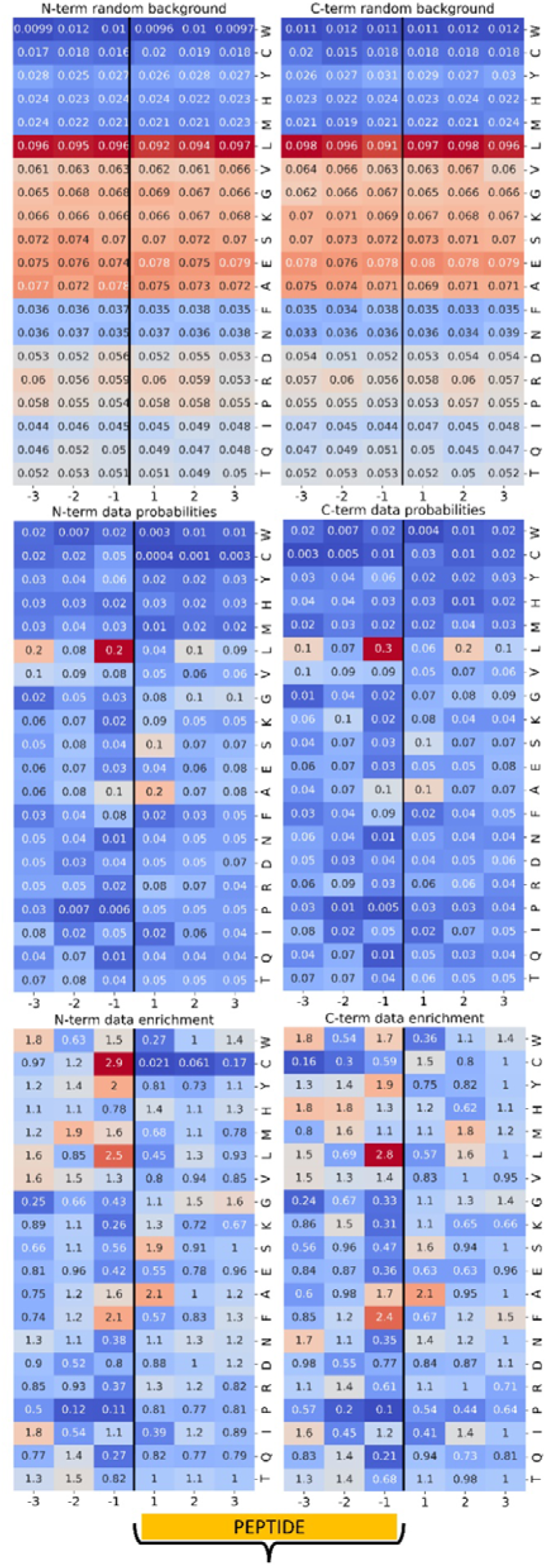
Amino acid enrichment in regions flanking N- and C- termini of intracellular peptides detected in A374 melanoma cell line. The topmost row shows a distribution of randomly selected peptides as a comparison. The middle row shows actual AA frequencies around the termini of detected peptides. The bottom row shows an enrichment factor calculated by dividing actual and random AA frequencies.

### Obtained peptide fraction reflects the native intracellular peptidome population

It can be speculated that peptides isolated using the methodology described here can be primarily a product of artifactual protein degradation during lysis. To ensure a predominantly native character of these peptides, we have analyzed their population behavior in conditions where it can be at least partially predicted. In a subsequent experiment, we compared the intracellular peptidome of SUP-T1 cells before and after the inhibition of proteasome activity by bortezomib. This drug binds to two of three proteasome subunits, affecting predominantly chymotrypsin-like activity. Trypsin-like activity should remain unchanged. Subunit activity measurements^40^ (Figure 10A) confirmed the predicted inhibition pattern, and the peptidome exhibited high-magnitude changes (Figure 10B). A closer examination of the dependency of abundance change on peptide cut type revealed that chymotryptic peptides are predominantly downregulated, whereas tryptic are mostly unaffected (Figure 10C).

**Figure 10.**
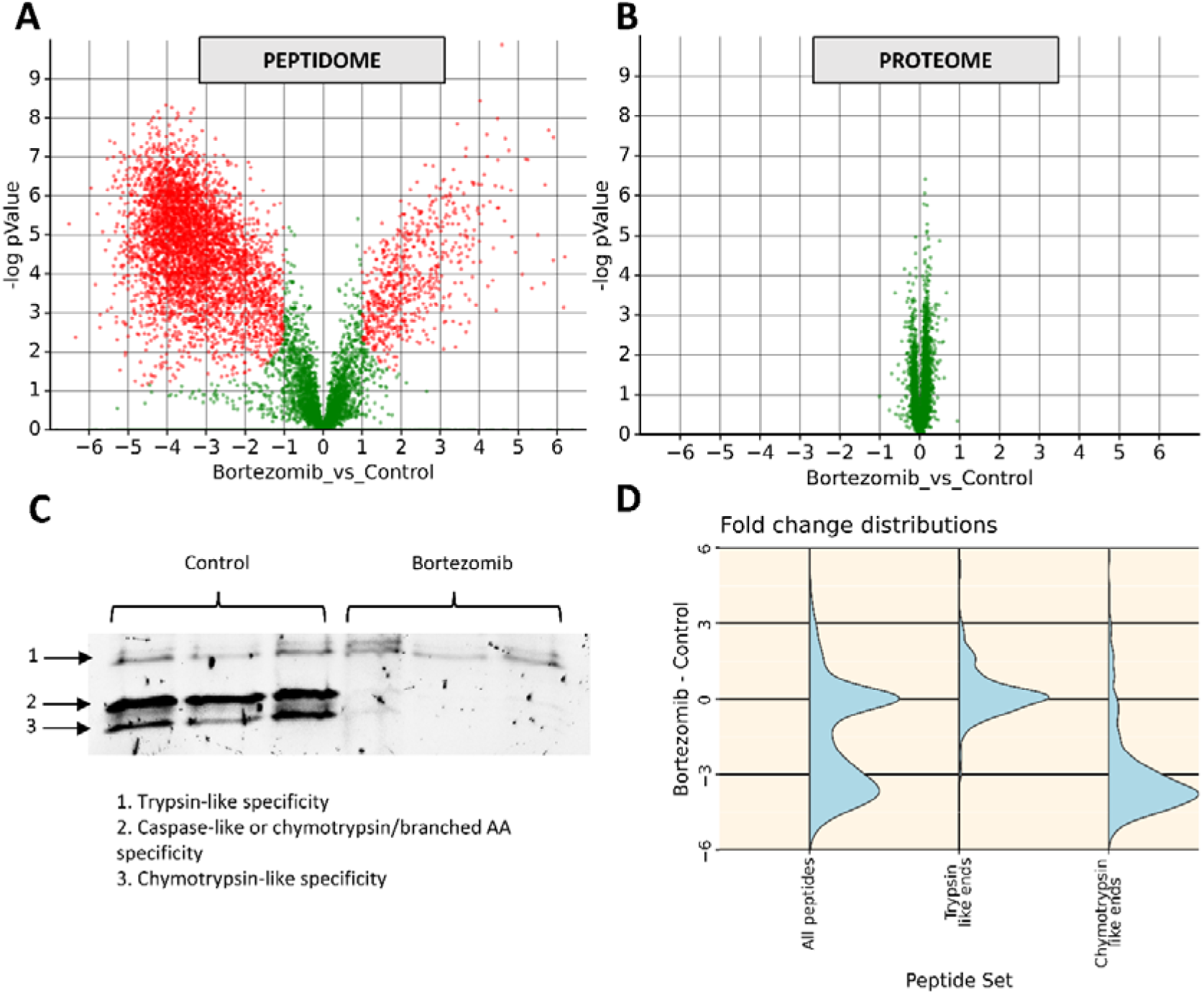
Quantitative peptidome analysis is applied to investigate the effects of proteasome inhibition. A) Volcano plot showing peptide fold changes after proteasome inhibition. B) The volcano plot shows protein fold changes after proteasome inhibition. C) Measurement of proteasome activity inhibition by bortezomib. Proteasome subunits were labeled with activity-based probes in 3 technical repeats (see Methods section). D) Comparison of fold change distribution of peptide produced with different cut sites.

These results show that measured changes in peptide abundances reflect the expected behavior of the proteasome product population. This allows us to confirm that we can analyze the native intracellular peptide population and get realistic insights into its behavior.

### In-depth peptidome/small proteome characterization by direct peptide fractionation

Knowing that HLB SPE results in a strong enrichment of small proteins and peptides even without the ultrafiltration step (Supplementary Figure 5), we leveraged it for comprehensive analysis of small proteome fraction isolated from the A549 non-small cell lung cancer (NSCLC) cell line. We have performed a comprehensive analysis of fractionated intact polypeptide filtrates and the identification of tryptic peptides obtained after the digestion of these fractions. Additionally, we assessed how the acquired data, combined with the previously acquired total proteome (fractionated tryptic digest) dataset, can be applied to identify novel proteoforms derived from small open reading frames (sORFs). For this purpose, we utilized a search database maintained at sorfs.org^41^, containing sequences predicted using ribosomal profiling data. As the retention character of HLB resin differs from typical C18 silica-based materials, we have used these columns directly to fractionate them. We have tested three fractionation conditions, namely, low pH methanol (MeOH), low pH acetonitrile (FA), and high pH acetonitrile (TAE), as described in detail in the Methods section.

The peptide mass range was extended above 10 kDa but did not contain intact proteins bigger than 20 kDa (Figure 11B). Obtained data posed a significant challenge during analysis, as search tools designed for bottom-up proteomics cannot handle complex spectra from long peptides. Therefore, we have combined search results obtained with the MSFragger search engine designed for bottom-up experiments with results obtained using MSPathFinder – a tool designed for top-down protein characterization.

**Figure 11.**
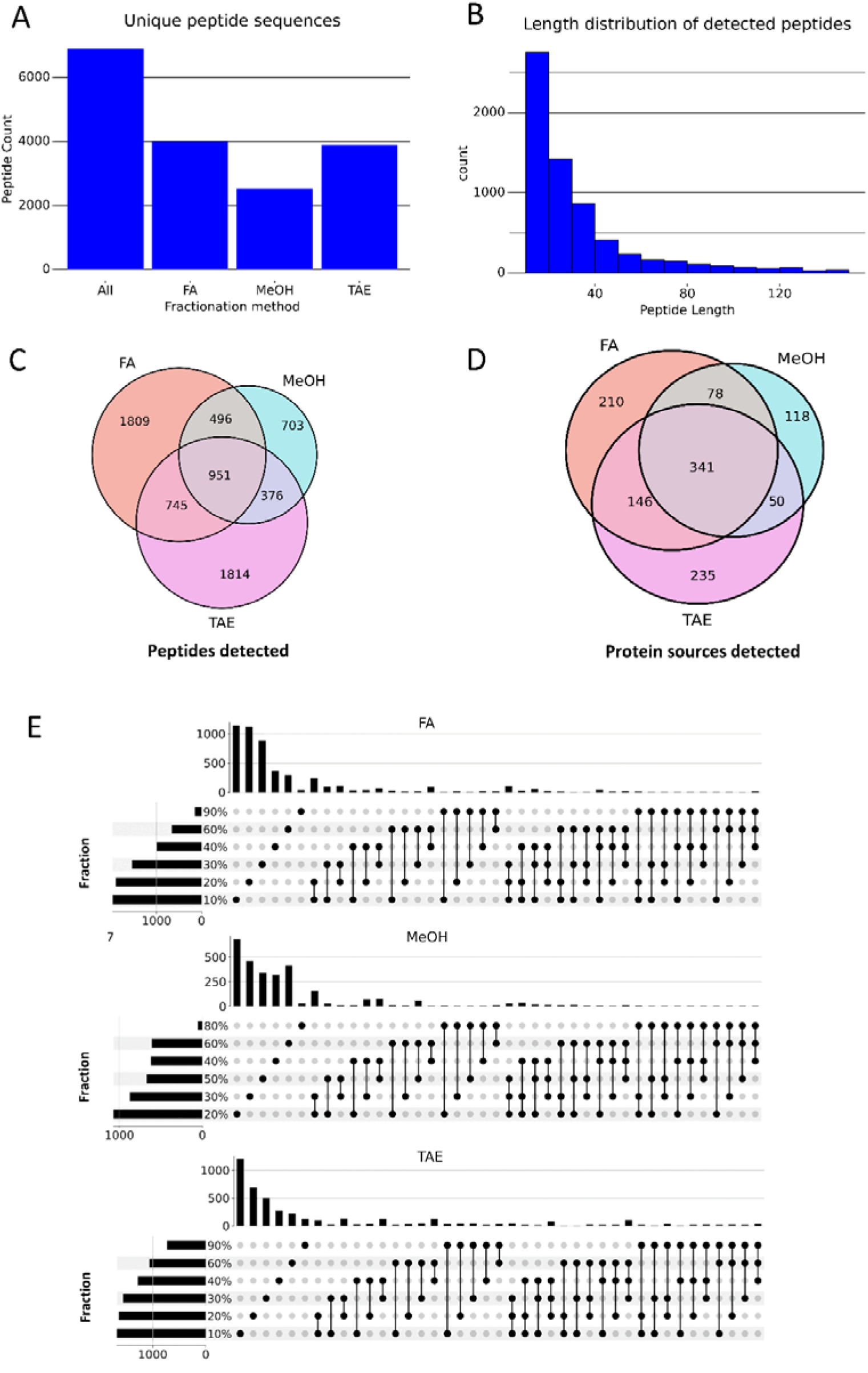
Results of peptidome/small proteome fractionation using different elution methods. A) Number of unique peptide sequences detected using different fractionation methods. B) The length distribution of peptides detected in all fractionation methods is combined. C) Venn diagram of peptide detections (unique sequences) for tested fractionation methods. D) Venn diagram of possible protein sources of detected peptides for tested fractionation methods. E) Upset plots showing fractionation efficiency of tested methods.

The relatively small number of identified peptides in fractionated samples compared to samples depleted of longer peptides (Figure 11A, compared with 16076 peptides quantified in melanoma experiment) suggests a substantial signal suppression coming long peptides and abundant components of the sample. This shows that size exclusion is beneficial if the experiment focuses on identifying or quantifying shorter peptides. Venn diagrams (Figure 11C, D) comparing the intracellular peptides identified in different fractionation conditions suggest that the eluted peptides provide different peptide populations. These observations reveal the difference in elution mechanisms from the HLB column, depending on the buffer used. Importantly, Figure 11E clearly shows that all compared elution buffers effectively separate vast portions of peptides into single or two adjacent fractions. Fractionation is also sufficiently orthogonal to separation on the C18 column.

We have performed identical sample analysis after trypsin digestion to check the extent of recovered and analyzed sequence diversity. Utilizing trypsin digestion leads to losing information about the intact peptide sequences. However, it provided considerably more data on possible protein sources of peptides of interest, as shown in Figure 12. The vast difference in the number of possible protein sources detected suggests that the detection of longer polypeptides is suboptimal and requires improvement.

**Figure 12.**
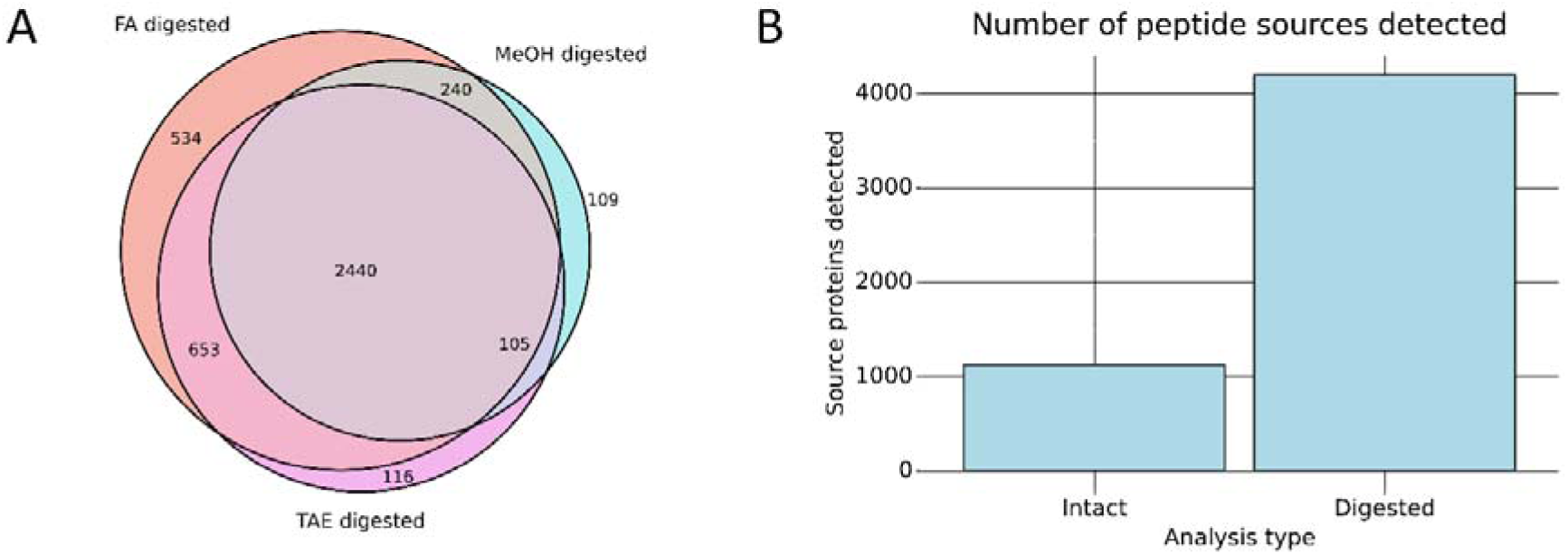
Analyses of trypsin-digested small proteome fractions. A) Venn diagram showing overlap of possible intracellular peptide sources detected through fractionation methods after tryptic digestion of fractions. B) Improve detection of possible peptide sources after tryptic digestion of fractions compared to undigested (intact) fractions.

We have combined peptide identifications from intact and digested fractions with those from high pH reverse-phase fractionated A549 proteome digest to detect potential sORFs (Figure 13). Furthermore, we have performed additional searches of intact fractions, using a smaller database containing all possible protein sources detected to enhance further peptide identification (see Methods section for details). Overall, peptide identification efficiency was only moderately improved, and novel possible protein sources were mostly derived only from single peptide identification. However, this approach led to the detection of an additional 6 sORF products on at least two unique peptides (Figure 13C). Results from sORF identification are summarized in Figure 13. All sORF products detected on minimum 2 peptides are shown in Supplementary Table 2.

**Figure 13.**
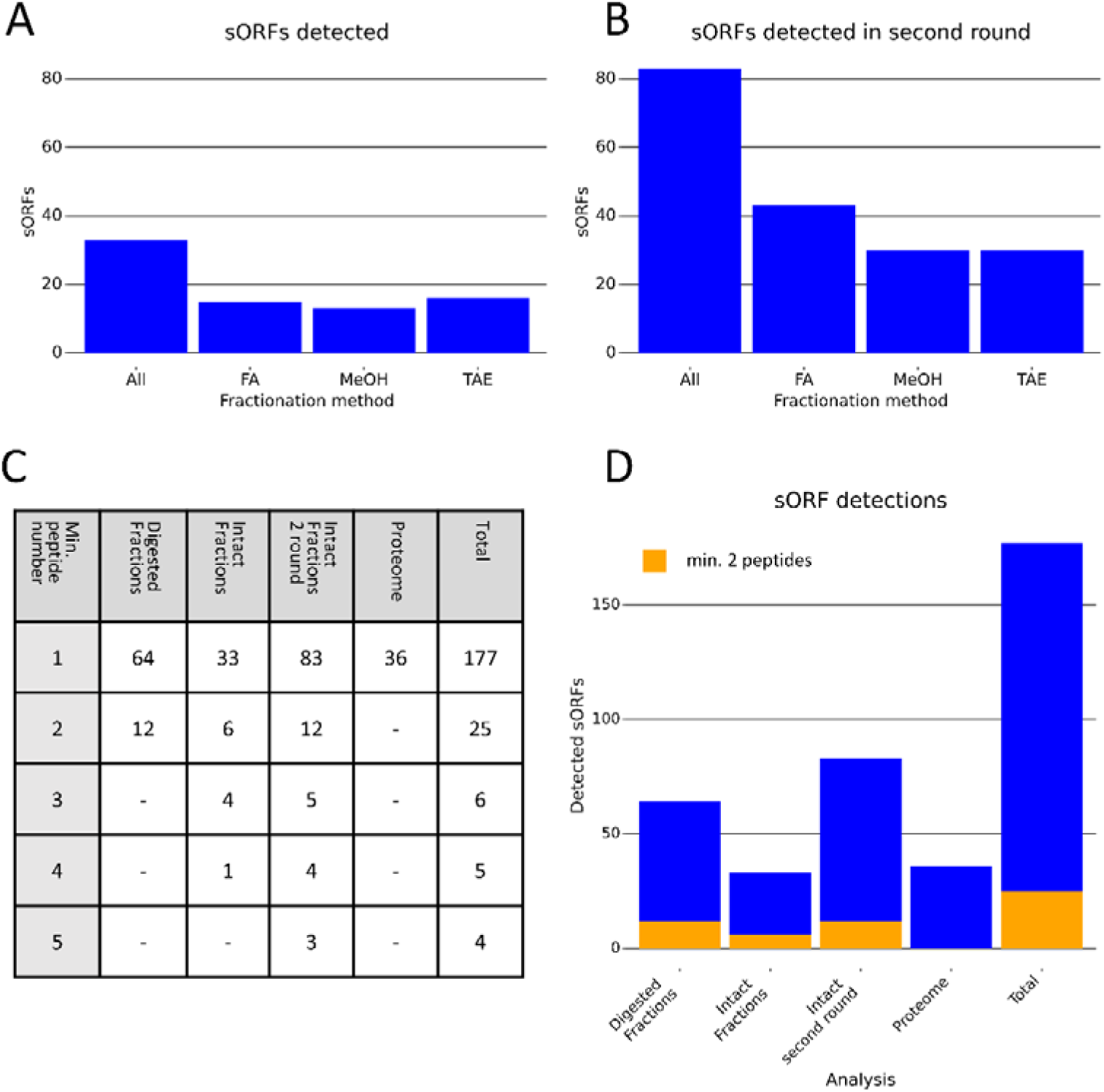
Detection of non-canonical sORF products using combined data. A, B) Comparison of a number of potential sORFs detected without (A) and with (B) additional search rounds on the reduced database for different fractionation methods. C) Number of potential sORFs detected with at least n peptides in different analyses: Intact Fractions - dataset from intact fraction runs of all fractionation methods, standard database, Digested Fractions - dataset from trypsin digested fraction runs of all fractionation methods, standard database, Intact Fractions 2 round - dataset from intact fraction runs of all fractionation methods, reduced database, Proteome - data from high-pH fractionated A549 proteome digest, standard database. D) Graphical representation of data from Table C for all potential sORF candidates and candidates detected with at least 2 peptides.

The number of sORFs and peptides detected per sORF were calculated exclusively using peptides not present in the Uniprot database and by generating a minimal sORF set that can explain the existence of peptides detected in experimental data. The number of sORF detections is similar to previously reported attempts^8,27^, despite using a generic sORF sequence database rather than a database generated by ribosome profiling of the analyzed cell line, which might result in better coverage.

## Discussion and future perspectives

In our hands, the methods provided here started to serve as valuable methods for characterizing intracellular peptidome and small proteome. We wish to provide a framework for replicating and developing intracellular peptidomics according to the researchers’ needs. One of the substantial aspects of the provided method is compatibility with a wide range of lysis conditions. It is important to highlight that it is well-applicable for all downstream intracellular peptidomics steps if a lysis solution can be diluted to have a less than 5% organic content. However, caution must be taken to avoid compounds that may be retained and eluted together with peptides. For instance, excessive amounts of peptide-based protease inhibitors may obscure signals of peptide ions. This sets the stage for new experiments aiming to characterize the efficiency of peptide extraction and deactivation of intracellular proteases using a wider range of conditions.

As most of the studies described in the literature apply ultrafiltration devices to remove the bulk of the high-mass compounds directly after lysis^17,20^, we have initially tested the feasibility of this method. Nevertheless, in our hands, it proved to be a very inefficient approach, leading to almost complete sample loss in most of the cases. Therefore, we recommend clearing the sample with SPE, which provides a much better-starting material for later ultrafiltration. Otherwise, the filter might become clogged by protein aggregates and/or other large biomolecules. It should be noted that even with this preparation, low molecular cut-off filters do not exhibit the anticipated level of effectiveness (refer to Figure 4B). Consequently, exploring an alternative method that retains the simplicity of ultrafiltration would be advantageous if one would like to push the limit of minimal sample amount requirements.

Intracellular peptide quantification provided surprisingly good results using the optimized intracellular peptidomics protocol. We maintained the simplicity of sample preparation and data analysis by using DIA label-free approach and freely available analytical software. The simplicity and comprehensiveness ensure that anyone with access to LC/MS systems of similar quality can quickly implement this novel method. The wide availability of the workflow may lead to a complete annotation of intracellular peptides and their functions in future studies, which is a prerequisite for the development of the peptidomics field. Analysis of peptides in length range >50 amino acids, which fall into the ‘middle-down’ analysis category, poses a much bigger challenge. Nevertheless, a middle-down approach for intracellular peptidomics will be necessary to adequately characterize novel proteoforms, which cannot be distinguished from full-length proteins after proteolysis.

We have shown that a small molecular weight fraction of the proteome is characterized by great diversity. Its characterization will undoubtedly benefit from more advanced computational and technical approaches. Using complementary fragmentation techniques such as ETD, ECD or EAD, whenever available, would be an advantage in generating more interpretable fragmentation spectra of large peptides. Developing workflows designed to ‘middle-down’ peptide/protein identification, either using intact peptide/protein data alone or with data from proteolytically digested samples, will make an analysis substantially easier. Some of such tools are being developed^42,43^, however they are less mature than more conventional software for bottom-up data analysis. Optimization of liquid chromatography conditions may also lead to an increase in the data quality. In the current manuscript, we utilized 300A C18 column material, which is relatively universal for separating a wide mass range peptide. However, using more specialized conditions and column materials (such as a monolithic polymer) may be beneficial. Unfortunately, such column chemistry was not readily available in nano-LC format when performing analyses. Moreover, the unique features of our novel peptidomics method compared to existing peptidomics and proteomics methods are in Supplementary Table 3. Through its application, we have shown the multipurpose nature of intracellular peptidomics, and a possible wide range of basic and applied science applications are visualized in Figure 1. Further application of our work can provide biologically relevant insights into various aspects of peptidome analysis, including active peptide biomarker discovery, identifying sORF products, and protein degradation analysis. In summary, our focus has been on providing a novel, simple, cost-effective, comprehensive, and globally applicable tool for the research community interested in peptidome/small proteome characterization that can be easily adapted and developed further

## Materials and methods

For clarity, a detailed description of all experimental procedures is available in supplementary material.

### Cell lysis for peptidome extraction recommended for quantitative analysis

Frozen pellets of 2×10^7^ cells were swiftly resuspended and vigorously pipetted in 1 ml of 8 M urea dissolved in PBS with 1 mM of PMSF and 1mM EDTA. Subsequently, samples were vortexed for 30 min and sonicated for 30 min at room temperature. Cellular debris were removed by centrifugation for 20 min at 17 kG. Supernatant was diluted to 6 ml with 0.2% formic acid. Polypeptides retained on the cartridge were three times washed with 1 ml of 5% MeOH, 0.2% FA in water.

### HLB-based sample cleanup

During all steps, cartridges were operated exclusively by gravity flow. HLB cartridges (30 mg; Waters, hydrophilic-lipophilic balanced columns, P/N: 186000382) was first conditioned with 1 ml of 0.2% FA in methanol (MeOH) and equilibrated with 1 ml of 0.2% FA. Diluted lysates were additionally centrifuged for 10 min at 5 kG to remove residual debris and aggregates, and immediately loaded on the cartridges. Polypeptides retained on the cartridge were three times washed with 1 ml of 5% MeOH, 0.2% FA in water. Finally, bound material was eluted with 1ml of 80% MeOH, 0.2% FA. Eluates were diluted to 40% MeOH, 0.2% FA in water with 0.2% FA in water prior filtration step, to ensure compatibility with regenerated cellulose membranes.

Above mentioned optimized basic workflow (lysis, cleanup and ultrafiltration) was applied for peptidome isolation for analysis of the effect of bortezomib.

### Data analysis – peptide identification

For shorter peptides we have used FragPipe suite equipped with MSFragger^44^ search engine. It was reasonable choice for searching large sequence spaces encountered in peptidome identification. As described in Table 5, it was routinely possible to search nonspecific peptides of length range 7 to 60 amino acids within complete human reference proteome (all isoforms) even when concatenated with additional sORF database. To test the possibilities of identifying even longer polypeptides, we have used MSPathfinderT tool included in Informed Proteomics^45^ package, which is intended for top-down protein analysis. It enabled for identifying small proteoforms up to approximately 15kDa. Using this tool, it is necessary to limit the search space (e.g to reviewed Swissprot sequences), as analysis time could easily exceed 24 hours per 120 min MS run on 32 core CPU.

**Table 5.** Summary of peptide search methods used in described experiments. UP AI = all isoformsUP = Uniprot; SP = SwissProt; TR = TrEMBL; RDB = reduced database all proteins detected 44467 entries; IP = Informed Proteomics, FRG = MSFragger, All searches were performed with following variable modifications: Methionine oxidation, N-term acetylation.

| Experiment | Method no. | File conversion , format | Search engine | Search database | Search type | Precursor ion tolerance (ppm) | Product ion tolerance (ppm) | Peptide length/mass |
| --- | --- | --- | --- | --- | --- | --- | --- | --- |
| Lysis buffer comparison and cut-off filter comparison |  | MSConvert .mzML | FRGv3.7 | UP:SP+TR AI, DI | Nonspecific | 10 | Auto | 7-60 |
| Intact peptide search for fractionation | 1 | MSConvert .mzML | FRG v3.7 | UP:SP+TR AI | Nonspecific | 10 | Auto | 7-40 |
|  | 2 | MSConvert .mzML | FRG v3.7 | UP:SP+TR AI | Nonspecific | 10 | Auto | 30-80 |
|  | 3 | MSConvert .mzML | FRG v3.7 | UP:SP+TR AI +sORF | Nonspecific | 10 | Auto | 7-40 |
|  | 4 | MSConvert .mzML | FRG v3.7 | UP:SP+TR AI +sORF | Nonspecific | 10 | Auto | 30-80 |
|  | 5 | PbfGen, .pbf | MSPathFinderT | UP:SP AI | Nonspecific | 8 | 16 | 3000-30000 |
|  | 6 | PbfGen, .pbf | MSPathFinderT | UP:SP AI +sORF | Nonspecific | 8 | 16 | 3000-30000 |
| Digested fractionated peptides | 1 | MSConvert .mzML | FRG v3.7 | UP:SP+TR AI | Semitryptic | 8 | Auto | 7-40 |
|  | 2 | MSConvert .mzML | FRG v3.7 | UP:SP+TR AI | Non-specific | 8 | Auto | 7-40 |
|  | 3 | MSConvert .mzML | FRG v3.7 | UP:SP+TR AI +sORF | Semitryptic | 8 | Auto | 7-40 |
|  | 4 | MSConvert .mzML | FRG v3.7 | UP:SP+TR AI +sORF | Non-specific | 8 | Auto | 7-40 |
| Intact peptide search for fractionation – round 2 | FRG | MSConvert .mzML | FRG v3.7 | RDB | Nonspecific | 10 | Auto | 7-60 |
|  | IP | PbfGen, .pbf | MSPathFinderT | RDB | Nonspecific | 8 | 16 | 3000-30000 |
| Melanona peptidome |  | MSConvert .mzML | FRG v3.5 | UP:SP+TR AI | Nonspecific | 10 | Auto | 7-60 |
| Bortezomib effect |  | MSConvert .mzML | FRG v3.8 | UP:SP+TR AI | Nonspecific | 10 | Auto | 7-60 |

### Data analysis – DIA based peptide quantification

DIA peptide quantifications were performed using DIA-NN^46^. Experiment-specific spectral libraries was prepared beforehand using EasyPQP tool supplied with FragPipe suite. For peptide quantification, MBR option was selected and precursor FDR was set to 1 %. Robust LC (high accuracy) option was selected. MS2 and MS1 mass accuracies, and scan window were set according to values recommended by DIA-NN. Then, results were preprocessed by diann-r R library, to obtain peptide-level quantification.

## Data availability

The mass spectrometry intracellular peptidomics data have been deposited to the Massive repository (ftp://MSV000096358@massive.ucsd.edu). Massive partner repository with the dataset identifier PXD050950.

## Authors’ contributions

A.P. methodology, experimental procedures, original draft manuscript writing preparation and review, J.F. data analysis, manuscript writing, A.D. cell culture experiments, N.M.T. funding acquisition, S.K. study conceptualization, supervision, manuscript original draft preparation, review and editing, funding acquisition.

## Acknowledgements

The authors would also like to thank the CI-TASK, Gdansk, and PL-Grid Infrastructure, Poland, for providing their hardware and software resources.

## Declarations

Competing interests. The authors declare no competing interests.

## Bibliography

1. Edwards, S. L. et al. Exploring neuropeptide signalling through proteomics and peptidomics. Expert Rev. Proteomics 16, 131–137 (2019).

2. Kim, J. S., Jeon, B. W. & Kim, J. Signaling Peptides Regulating Abiotic Stress Responses in Plants. Front. Plant Sci. 12, (2021).

3. Shapiro, I. E. & Bassani-Sternberg, M. The impact of immunopeptidomics: From basic research to clinical implementation. Semin. Immunol. 66, 101727 (2023).

4. Sandmann, C.-L. et al. Evolutionary origins and interactomes of human, young microproteins and small peptides translated from short open reading frames. Mol. Cell 83, 994–1011.e18 (2023).

5. Delcourt, V., Staskevicius, A., Salzet, M., Fournier, I. & Roucou, X. Small Proteins Encoded by Unannotated ORFs are Rising Stars of the Proteome, Confirming Shortcomings in Genome Annotations and Current Vision of an mRNA. PROTEOMICS 18, 1700058 (2018).

6. de Araujo, C. B. et al. Intracellular Peptides in Cell Biology and Pharmacology. Biomolecules 9, 150 (2019).

7. Morgan, G. R. & Carlyle, B. C. Interrogation of the human cortical peptidome uncovers cell-type specific signatures of cognitive resilience against Alzheimer’s disease. Sci. Rep. 14, 7161 (2024).

8. Cardon, T. et al. Optimized Sample Preparation Workflow for Improved Identification of Ghost Proteins. Anal. Chem. 92, 1122–1129 (2020).

9. Fabre, B., Combier, J.-P. & Plaza, S. Recent advances in mass spectrometry–based peptidomics workflows to identify short-open-reading-frame-encoded peptides and explore their functions. Curr. Opin. Chem. Biol. 60, 122–130 (2021).

10. Ferro, E. S., Rioli, V., Castro, L. M. & Fricker, L. D. Intracellular peptides: From discovery to function. EuPA Open Proteomics 3, 143–151 (2014).

11. Galindo, M. I., Pueyo, J. I., Fouix, S., Bishop, S. A. & Couso, J. P. Peptides Encoded by Short ORFs Control Development and Define a New Eukaryotic Gene Family. PLoS Biol. 5, e106 (2007).

12. Weinstein-Marom, H. et al. MHC-I presentation of peptides derived from intact protein products of the pioneer round of translation. FASEB J. 33, 11458–11468 (2019).

13. Becker, J. P. et al. Pharmacological inhibition of nonsense-mediated RNA decay augments HLA class I-mediated presentation of neoepitopes in MSI CRC. 2020.10.13.319970 Preprint at 10.1101/2020.10.13.319970 (2020).

14. Fricker, L. D. Analysis of mouse brain peptides using mass spectrometry-based peptidomics: implications for novel functions ranging from non-classical neuropeptides to microproteins. Mol. Biosyst. 6, 1355–1365 (2010).

15. Zhang, F. et al. Peptidome Analysis of Pancreatic Tissue Derived from T1DM Mice: Insights into the Pathogenesis and Clinical Treatments of T1DM. BioMed Res. Int. 2021, 9987042 (2021).

16. Dias, N. C. & Poole, C. F. Mechanistic study of the sorption properties of OASIS® HLB and its use in solid-phase extraction. Chromatographia 56, 269–275 (2002).

17. Zhou, C.-X. et al. Quantitative Peptidomics of Mouse Brain After Infection With Cyst-Forming Toxoplasma gondii. Front. Immunol. 12, (2021).

18. Zhang, L. et al. Peptidomics Analysis Reveals Peptide PDCryab1 Inhibits Doxorubicin-Induced Cardiotoxicity. Oxid. Med. Cell. Longev. 2020, 7182428 (2020).

19. Wu, L. et al. Peptidomic Analysis of Cultured Cardiomyocytes Exposed to Acute Ischemic-Hypoxia. Cell. Physiol. Biochem. 41, 358–368 (2017).

20. Li, X. et al. Comparative peptidome profiling reveals critical roles for peptides in the pathology of pancreatic cancer. Int. J. Biochem. Cell Biol. 120, 105687 (2020).

21. Xue, Y. et al. Peptidomic Analysis of Endometrial Tissue from Patients with Ovarian Endometriosis. Cell. Physiol. Biochem. 47, 107–118 (2018).

22. Sahu, I. et al. The 20S as a stand-alone proteasome in cells can degrade the ubiquitin tag. Nat. Commun. 12, 6173 (2021).

23. Zhang, J. et al. Peptidome analysis reveals critical roles for peptides in a rat model of intestinal ischemia/reperfusion injury. Aging 15, 12852 (2023).

24. Dasgupta, S. et al. Proteasome Inhibitors Alter Levels of Intracellular Peptides in HEK293T and SH-SY5Y Cells. PLOS ONE 9, e103604 (2014).

25. Li, Y. et al. Identification of intracellular peptides associated with thermogenesis in human brown adipocytes. J. Cell. Physiol. 234, 7104–7114 (2019).

26. Parmar, B. S. et al. Identification of Non-Canonical Translation Products in C. elegans Using Tandem Mass Spectrometry. Front. Genet. 12, (2021).

27. Ma, J. et al. Discovery of Human sORF-Encoded Polypeptides (SEPs) in Cell Lines and Tissue. J. Proteome Res. 13, 1757–1765 (2014).

28. Kawashima, Y. et al. Optimization of Data-Independent Acquisition Mass Spectrometry for Deep and Highly Sensitive Proteomic Analysis. Int. J. Mol. Sci. 20, 5932 (2019).

29. Matsumoto, G., Wada, K., Okuno, M., Kurosawa, M. & Nukina, N. Serine 403 phosphorylation of p62/SQSTM1 regulates selective autophagic clearance of ubiquitinated proteins. Mol. Cell 44, 279–289 (2011).

30. Suzuki, N. et al. TP53/p53-FBXO22-TFEB controls basal autophagy to govern hormesis. Autophagy 17, 3776–3793 (2021).

31. Zhang, Y. F. et al. Nuclear localization of Y-box factor YB1 requires wild-type p53. Oncogene 22, 2782–2794 (2003).

32. Katte, R. H., Chou, R.-H. & Yu, C. Pentamidine inhibit S100A4 - p53 interaction and decreases cell proliferation activity. Arch. Biochem. Biophys. 691, 108442 (2020).

33. Li, Z.-H. & Bresnick, A. R. The S100A4 metastasis factor regulates cellular motility via a direct interaction with myosin-IIA. Cancer Res. 66, 5173–5180 (2006).

34. Tag7-Mts1 Complex Induces Lymphocytes Migration via CCR5 and CXCR3 Receptors - PubMed. https://pubmed.ncbi.nlm.nih.gov/30713770/.

35. Proteoform Atlas – Consortium for Top Down Proteomics. http://atlas.topdownproteomics.org/proteoforms/18188.

36. Dasgupta, S. et al. Analysis of the Yeast Peptidome and Comparison with the Human Peptidome. PLOS ONE 11, e0163312 (2016).

37. Teixeira, C. M. M., Correa, C. N., Iwai, L. K., Ferro, E. S. & Castro, L. M. de. Characterization of Intracellular Peptides from Zebrafish (Danio rerio) Brain. Zebrafish 16, 240–251 (2019).

38. Lyapina, I., Ivanov, V. & Fesenko, I. Peptidome: Chaos or Inevitability. Int. J. Mol. Sci. 22, 13128 (2021).

39. Wolf-Levy, H. et al. Revealing the Cellular Degradome by Mass Spectrometry Analysis of Proteasome-Cleaved Peptides. Nat. Biotechnol. 10.1038/nbt.4279 (2018) doi:10.1038/nbt.4279.

40. de Bruin, G. et al. A Set of Activity-Based Probes to Visualize Human (Immuno)proteasome Activities. Angew. Chem. Int. Ed Engl. 55, 4199–4203 (2016).

41. Olexiouk, V., Van Criekinge, W. & Menschaert, G. An update on sORFs.org: a repository of small ORFs identified by ribosome profiling. Nucleic Acids Res. 46, D497–D502 (2018).

42. Lima, D. B. et al. ProteoCombiner: integrating bottom-up with top-down proteomics data for improved proteoform assessment. Bioinforma. Oxf. Engl. 37, 2206–2208 (2021).

43. Schaffer, L. V., Millikin, R. J., Shortreed, M. R., Scalf, M. & Smith, L. M. Improving Proteoform Identifications in Complex Systems Through Integration of Bottom-Up and Top-Down Data. J. Proteome Res. 19, 3510–3517 (2020).

44. Kong, A. T., Leprevost, F. V., Avtonomov, D. M., Mellacheruvu, D. & Nesvizhskii, A. I. MSFragger: ultrafast and comprehensive peptide identification in mass spectrometry–based proteomics. Nat. Methods 14, 513–520 (2017).

45. Park, J. et al. Informed-Proteomics: Open Source Software Package for Top-down Proteomics. Nat. Methods 14, 909–914 (2017).

46. Demichev, V., Messner, C. B., Vernardis, S. I., Lilley, K. S. & Ralser, M. DIA-NN: neural networks and interference correction enable deep proteome coverage in high throughput. Nat. Methods 17, 41–44 (2020).

